# Live single molecule microscopy of HIV-1 assembly in host T cells reveals a spatio-temporal effect of the viral genome

**DOI:** 10.1101/267930

**Authors:** Charlotte Floderer, Jean-Baptiste Masson, Elise Boiley, Sonia Georgeault, Peggy Merida, Mohamed El Beheiry, Maxime Dahan, Philippe Roigeard, Jean-Baptiste Sibarita, Cyril Favard, Delphine Muriaux

## Abstract

Monitoring virus assembly dynamic at the nanoscale level in host cells remains a major challenge. Human Immunodeficiency Virus type 1 (HIV-1) components are addressed to the plasma membrane where they assemble to form spherical particles of 100nm in diameter. HIV-1 Gag protein expression alone is sufficient to produce virus-like particles (VLPs) that resemble immature virus. Here, we monitored Gag assembly in host CD4 T lymphocytes using single molecule dynamics microscopy and energy mapping. A workflow allowing long time recordings of single Gag molecule localization, diffusion and effective energy maps was developed for robust quantitative analysis of HIV assembly and budding. Comparison of numerous cell plasma membrane assembling platforms in cells expressing wild type or assembly-defective Gag proteins showed that VLP formation last 15 minutes, with an assembly time of 5 minutes, and that the nucleocapsid domain is mandatory. Importantly, it reveals that the viral genome coordinates spatio-temporally HIV-1 assembly.

## Introduction

Studying enveloped RNA virus assembly at the host cell surface is crucial for understanding the specificities and differences of the underlying molecular mechanisms. However, this requires tools to enable the nanoscale analysis of viral proteins at the single molecule level. For instance, human immunodeficiency virus type 1 (HIV-1) produces particles with a diameter of 100-130nm filled with 2000 Gag proteins. Recent progress in single-molecule localization microscopy allows deciphering protein organization and dynamics in a single cell at the nanoscale level (1, 2). HIV-1 Gag polyprotein is the main determinant for HIV-1 particle assembly that occurs mainly at the plasma membrane of the host cell (3). When expressed alone in a cell, HIV-1 Gag proteins can produce non-infectious virus-like particles (VLPs) that resemble viruses, but do not require maturation (encoded by the Pol gene) or envelope proteins (encode by the Env gene). Therefore, it is a very powerful tool for studying virus assembly in a minimal productive system (4). Upon virus maturation, HIV-1 Gag polyprotein is cleaved by the viral protease into the following domains: Matrix protein p17 (MA), Capsid protein p24 (CA), Nucleocapsid protein p7 (NC) as well as p6 domain and two spacer peptides (sp1 and sp2). MA is myristoylated and contains a highly basic region involved in Gag targeting and anchoring to the inner leaflet of the plasma membrane of host cells where viral assembly occurs (reviewed in (5–7). CA, via CA-CA interacting domains, promote Gag-Gag oligomerization in vitro (8, 9) and in cells (10). Particularly, the WM mutation within CA strongly reduces Gag oligomerization and consequently virus assembly (8). NC, sp2 and p6 are required for Gag assembly and particle budding. Specifically, NC recruits the genomic RNA, but can also interact with cellular RNAs, to favour Gag-Gag oligomerization on the RNA template, in vitro and in cells (reviewed in (11) and (12)). NC is therefore involved in virus assembly (12) and (13), whereas p6 recruits the ESCRT proteins required for membrane scission and particle release (14). The sp1, at the end of the CA, acts as a molecular switch for VLP assembly (15). The kinetics of GFP-labelled Gag assembly and VLP formation have been previously described in adherent HeLa cells by measuring the local increase in fluorescent intensity of single virions (16, 17). In these cells, it was estimated that 5 to 6 minutes were required for Gag VLP assembly in the absence of genomic RNA (16) and about 20 minutes to complete 90% of Gag VLP assembly and budding (17). Jouvenet *et al.* (18) also reported that HIV-1 Gag and a fluorescent tagged viral RNA assemble at the plasma membrane. Moreover, it was recently shown that HIV-1 Gag assembly at the plasma membrane takes place at sites where the viral RNA is located (19). The viral Gag proteins appears to stabilize the viral RNA at the plasma membrane and between 1/10 and 1/3 of the viral RNA is packaged into nascent particles in 30 minutes (20). These interactions between the viral genome and Gag enhance virus assembly (21). It suggested that the viral genomic RNA encoding Gag acts as a catalyser for virus assembly but it did not assess quantitatively the effect of the viral genomic RNA on the spatio-temporal coordination of HIV-1 Gag assembly. In this study, we wanted to measure HIV-1 Gag proteins dynamic changes during its assembly into a VLP, at the inner living cell surface, and to quantify the role of the different Gag domains, and of the viral RNA, upon VLP formation in the host CD4 T lymphocytes. To this aim, the photoactivable fluorescent tag mEOS2 was introduced into the HIV-1 Gag precursor (Gag(i)mEOS2). This tag allows using live photoactivated localization microscopy (PALM) (1, 22). Indeed, live PALM provides a precise spatio-temporal description of VLP formation, at the surface of individual cells, with a spatial resolution of about 50 nanometres and temporal resolution in the millisecond range. By coupling live PALM, TIRF-microscopy, and statistical analyses based on millions of Gag protein localizations and hundreds of buds, we could monitor non-infectious immature HIV-1 VLP formation at the cell plasma membrane of CD4 T cells molecules after molecules. Moreover, by comparing wild type (WT) and assembly-defective Gag mutants, we could identify which Gag domains are crucial for Gag assembly coordination at the host T cell surface. First, we found that in fixed CD4 T cells, Gag assembly platforms are rarely formed at the cell surface when Gag C-terminal end is deleted (the portion that contains the NC-viral RNA interaction domain). Then, based on the temporal changes observed in localization density maps, we showed that Gag VLP assembly in T cells requires between 5 and 7 minutes and 15min total to complete budding. Finally, by combining live PALM Bayesian inference analysis of single protein dynamic interaction maps with a diffusion and effective energy trapping model (23, 24), we quantified Gag trapping energy during assembly. Moreover, we analysed the temporal correlation between changes in the density and the trapping energy, ie Gag interaction, during VLP assembly and brought evidence that the cis-packageable viral genome that encodes Gag(i)mEOS2 spatio-temporally most probably coordinates VLP assembly at the cell surface of CD4 T lymphocytes.

## Results

### Expression of WT Gag(i)mEOS and assembly-defective mutants in Jurkat T cells

First, WT HIV-1 Gag(i)mEOS2 and known assembly defective mutants that harbour the mEOS2 tag were generated (Fig. 1a). The tag was introduced between MA and CA, thus preserving Gag capacity to assemble and to bud from the cell membrane after transient expression in mammalian cells (25). WT Gag(i)mEOS2 was produced using either the pNL4.3ΔPolΔEnv plasmid that includes also a cis-Psi-signal on the viral RNA that promotes viral RNA packaging into the VLP (NL4.3ΔPolΔEnv Gag, here after) (26) or the pGag(i)mEOS2 WT plasmid without this signal (WT Gag, hereafter). The WM, MACASP1, MACASP1/WM and Δdp6 Gag mutants were derived from WT Gag(i)mEOS2 (see Methods). WM harbours a mutation in CA that reduces CA-CA interactions and impairs Gag oligomerization. The MACASP1 mutant carries a stop codon at the end of CA-SP1 and therefore, lacks Gag C-terminus (NC-sp1-p6) (27). In Δp6, a deletion in the p6 domain of Gag impairs ESCRT recruitment and consequently particle release (14). Indeed, tethered Δp6 particles remained attached to the cell membrane (Supplementary Fig. S1a), as previously reported (14). All the Gag proteins were well expressed after transient transfection in Jurkat CD4 T lymphocytes, as indicated by western blot analysis (Fig. 1b) and by flow cytometry (Fig. 1e). Cell viability analysis by flow cytometry indicated that 40-50% of cells were alive after electroporation (Fig. 1d). Analysis of the geometric mean of fluorescent intensity (Fig. 1e) showed that 24 hours post-transfection, the global protein expression level was comparable for WT Gag and NL4.3ΔPolΔEnvGag, whereas it was 2-fold lower, on average, for MACASP1, WM and mEOS2 (vector alone). Finally, VLP production was assessed by semi-quantitative western blot analysis with an anti-CA antibody (Methods) (Fig. 1b and 1c). Only purified WT Gag and WM VLPs could be easily observed, whereas MACASP1 VLPs were often undetectable (Fig. 1b). In agreement, VLP release calculation showed that VLP production decreased from about 50% to 10% for MACASP1 (Fig. 1c). The capacity of WT Gag and mutants to bind to cell membranes was checked with membrane flotation assays in HEK293T cells (as described in Thomas *et al.* (27)) (Supplementary Fig. S2a). Although Gag was well expressed (Fig. 1b), the fraction of WT Gag bound to cell membranes was between 60 to 80% of total Gag (Supplementary Fig. S1b), and this value further decreased for WM and MACASP1 (p<0.01), and even more for the WM/MACASP1 double mutant (p<0.001). This made impossible analysis of the later by live PALM. These results indicate that upon alteration of Gag multimerization capacity, Gag is less bound to cell membranes and confirm a role for Gag oligomerization in stabilizing Gag-membrane interactions, in agreement with (28). Moreover, it was recently shown that in vitro Gag oligomerization occurs also on PIP2-containing lipid membranes and that it is reduced by the same WM mutation in the CA domain of Gag (9). Transmission electron microscopy was then used to check whether WT Gag(i)mEOS2 and mutants could form particles (Gag VLPs) (Supporting information). Upon expression of WT Gag, cells produced high amounts of electron-dense budding vesicles (i.e., Gag VLPs). After transfection of NL4.3ΔPolΔEnv Gag, particles seemed rarer at the cell surface than in WT Gag-expressing cells, possibly due to inefficient release. Quantification of the size of 40 to 150 particles (Supplementary Fig. S1b) indicated that the largest VLPs were produced in WM-expressing cells (205nm in diameter), whereas they were smaller (124nm in diameter on average) in WT Gag-expressing cells. Upon transfection of MACASP1, very rare VLPs were detected, often localized at the plasma membrane, as indicated by the dark staining at the cell membrane (Supplementary Fig. S1a). HEK293T cells can naturally produce some vesicles (mock) that are not electron-dense structures (Supplementary Fig. S1a). However, neither dark staining at the plasma membrane nor VLP was detected in mock cells (Supplementary Fig. S1a). These results are in agreement with the literature and allowed us to select the HIV-1 Gag protein variants that were well expressed in Jurkat T cells and that could bind to cell membranes, two prerequisites for analysing Gag(i)mEOS2 assembly at the cell surface by live PALM.

**Fig. 1.**
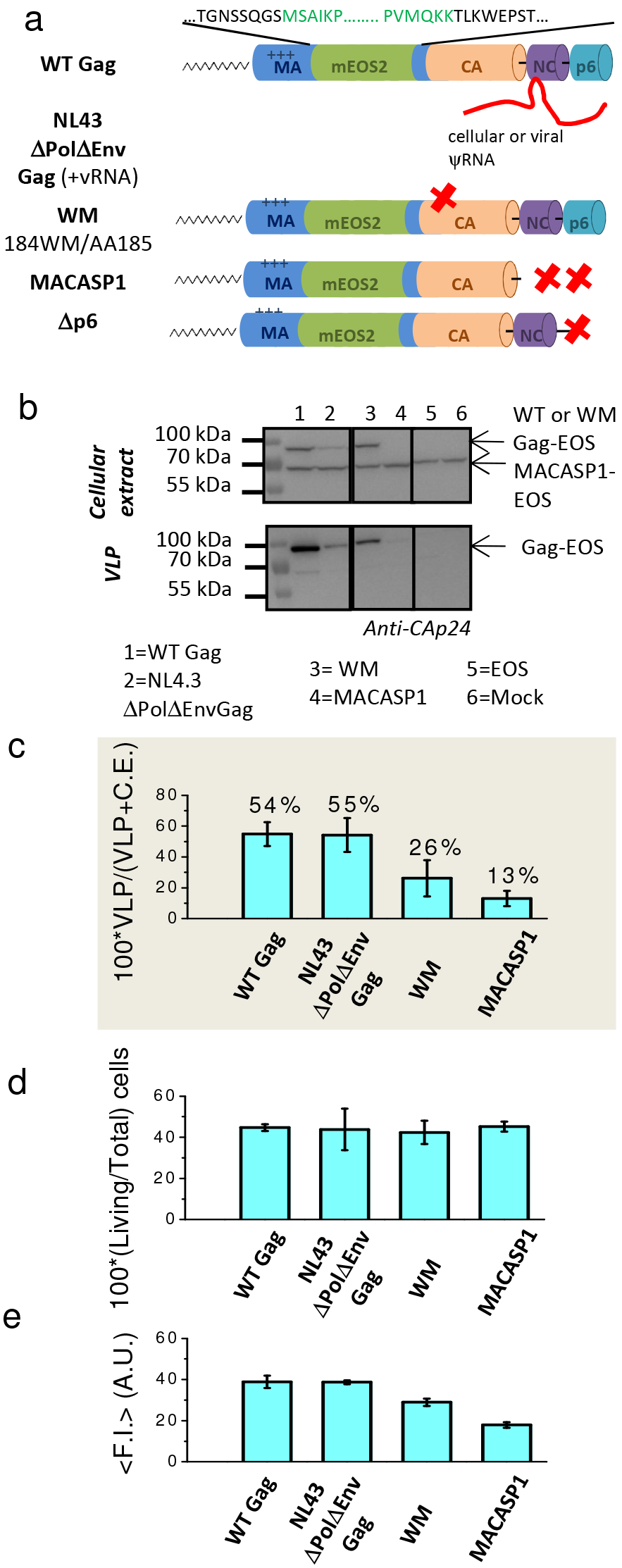
Characterization of Gag(i)mEOS2 wild type and mutants expressed in Jurkat T cells. (a) Schematic representation of HIV-1 WT Gag(i)mEOS2 and mutants used in this study, (b) Western blot analysis of HIV-1 WT protein and mutant expression in Jurkat T cells (“cellular extract”) and in purified VLPs (“VLP”). (c) Quantification of WT Gag(i)mEOS2 and mutant VLP release efficiency in transfected Jurkat T cells relative to total Gag protein (mean ± sd of 4 to 6 independent experiments). (d) Cell viability (relative to the total number of transfected cells) and protein expression of Gag(i)mEOS2 and mutants in Jurkat T cells using flow cytometry (mean ± sd of 3 independent experiments). (e) Global fluorescence intensity (F.I.) of Gag(i)mEOS2 transfected Jurkat T cells as measured by flow cytometry for each condition, as indicated.

### HIV-1 Gag mobility and assembly platform density are impaired at the Jurkat T cell surface upon deletion or mutation in Gag assembly domains

First, dynamics of single Gag molecules at the cell surface were analysed by reconstructing the trajectories from live PALM data acquired on CD4 Jurkat T cells in TIRF mode (Fig. 2a).

Using a simple diffusive model (Brownian motion), the instantaneous motion amplitudes were calculated by estimating the diffusion coefficient (D) from the mean square displacement (MSD) curve of each single-molecule trajectory. Although this measure is known to be noisy, the large number of short trajectories provided a valid estimate of the typical motion at the population scale. MSDs were computed for all trajectories longer than 8 frames obtained on the entire cell surface by single particle tracking PALM in 5-10 cells (see Methods). Then, all individual D values were pooled and their distribution was normalized and plotted on a logarithmic scale (Fig. 2b, open circles). Although these distributions exhibited slightly different maximal D values (from 0.11±0.12 *μm*^2^.*s*^−1^ for NL4.3ΔPolΔEnvGag to 0.35±0.4 *μm*^2^.s^−1^ for MACASP1), the high distribution variability did not allow us to easily identify the specific D features of Gag molecules in formed VLPs, in assembling platforms or diffusing as monomers. To overcome this issue, the D distribution of each Gag variant (Fig. 2b) were decomposed into different components using the D distributions of: i) CAAX(i)mEOS2 molecules anchored into the lipids of the T cell plasma membrane, to mimic the diffusion of Gag monomers at the beginning of the assembly, and ii) WT Gag(i)mEOS2 molecules irreversibly trapped in an already formed and released VLP, to mimic the diffusion at the assembly end (Fig. 2c). The maximal D was 0.6±0.5 *μm*^2^.*s*^−1^ for CAAX(i)mEOS2, and (2±0.8).10^−3^*μm*^2^.*s*^−1^ for trapped WT Gag(i)mEOS2 into a VLP (Fig. 2c). From this linear decomposition, an intermediate population could be extracted that was considered to represent the D distribution of Gag molecules in assembling platforms (green in Fig. 2b and d). These data showed that the immobile fraction (blue in Fig. 2b and d) was comparable for WT and mutant Gag molecules, whereas the intermediate and mobile fractions (green and red, respectively, in Fig. 2b and 2d) varied. The proportion of highly mobile WT proteins at the cell surface was very low (11 to 19% of all Gag molecules), while mutations in Gag oligomerization domains (WM for CA-CA interactions and MACASP1 for NC-RNA interactions) strongly increased this fraction to 39% and 52%, respectively. Consequently, the intermediate Gag population decreased from 82-69% for WT proteins to 54% and 34% for the WM and MACASP1 mutants, respectively. Again, this indicates that deleting Gag C-terminal domains, including the NC and p6, leads to an increase of highly mobile Gag fractions at the cell surface. This result is in good agreement with data on the measurement of GagΔNC mobility in HeLa cells (26, 29).

Then viral Gag clusters were observed in fixed Jurkat T cells. Examples of super-resolution reconstructed PALM images are presented in Fig. 3a for two different WT Gag (WT and NL43ΔPolΔEnv) and two different Gag oligomerisation-defective mutants (WM and MACASP1). Fig. 3b shows their associated distributions of the Gag clusters mean diameters and their log-normal fit obtained from several cells. Only MACASP1 is not following this fit. As shown in table1, the mean apparent diameters of WT Gag (112±54 nm) and NL4.3ΔPolΔEnv Gag (117±65nm) were not significantly different (see table S1 for statistics tests), but were significantly smaller than the WM one (121±56 nm). This was confirmed by electron microscopy (figure S1). Due to the low number of observed MACASP1 clusters, their mean diameter could not be determined. Finally, these mean diameter values were all found to be significantly smaller than the VLP one (139±42 nm) obtained from images of purified VLPs produced by WT Gag-expressing Jurkat T cells (Fig. 3a, WT purified VLPs), suggesting that theses clusters are Gag assembly platforms. The values obtained here are in agreement with previous PALM data on WT Gag assembly platform sizes described in adherent COS cells (1, 30) and suggests that the assembly platform size is independent of the host cell type. The PALM images were then used to quantify the platform density (i.e., the number of assembly platforms per cell surface units). It decreased from 4.5±1.7 for WT Gag and 3±1.2 clusters per *μm*^2^ for NL4.3ΔPolΔEnv Gag to 1.7± 1.5 clusters per *μm*^2^ for WM and to 0.05±0.01 for MACASP1 Gag mutants, suggesting that WM and MACASP1 increased mobility observed in Figure 2d partly reflects, at equilibrium, the decrease in assembling platform density observed at the CD4 T cell surface. Importantly, this result suggests a strong role of the Gag C-terminus domain (NC-sp2-p6) in assembly platform formation in CD4 T cells. Thus, PALM images of a Gag-Δp6 mutant (Fig. 3a and b, Δp6), which displays only the deletion of the p6 domain at the C terminus of Gag, were acquired. Δp6 mutant is a VLP release deficient mutant, forming grapes of VLPs (Inset of Fig. 3a and Supplementary Fig.S1a). This lead to an overestimation of the clusters mean diameter and underestimation of their density (Table1). Nevertheless, cells expressing Δp6 exhibited assembly platforms densities, and at least 20 times higher than the one observed for MACASP1 (Table 1), revealing that the NC domain of Gag was the major determinant for high density Gag clusters at the surface of Jurkat T cells.

**Fig. 2.**
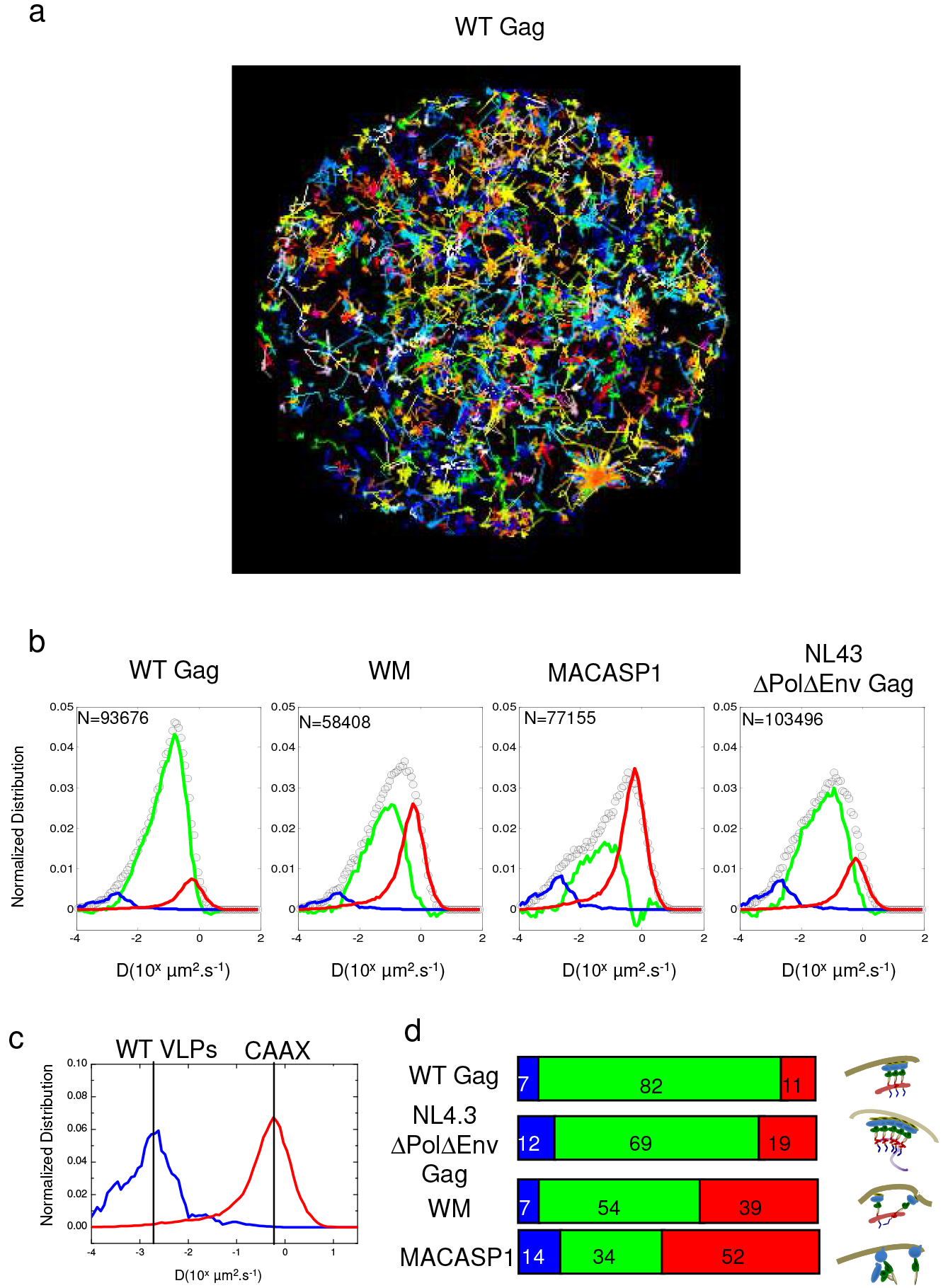
Gag(i)mEOS2 diffusion at the surface of Jurkat T cells and in VLPs. (a) Representative examples of the trajectories obtained in single cells for WT Gag and mutants. (b) Logarithmic distribution (open circles) of the diffusion coefficient D obtained in five cells for each con-dition. D values were obtained from the MSD linear fit of N trajectories (0.5.10^5^ <N<10^5^). (c) Logarithmic distribution of the diffusion coefficient D obtained for membrane-anchored CAAX(i)mEOS2 (control for molecules that do not assemble at the cell membrane) (red line) and VLPs (control for fully assembled and released particles) (blue line). These distributions were used for linear decomposition of the distribution plots in (b), and the remaining residual was plotted as the green curve in (b). Note that the apparent negative values in the distribution of D for the residual (green line) are a consequence of the linear decomposition of the obtained experimental curves (open circle). (d) Percentage of each com-ponent (assembled, assembling, monomers) observed for WT Gag and mutants. Each percentage corre-sponds to the weighted values of the linear decomposition obtained for each component.

**Fig. 3.**
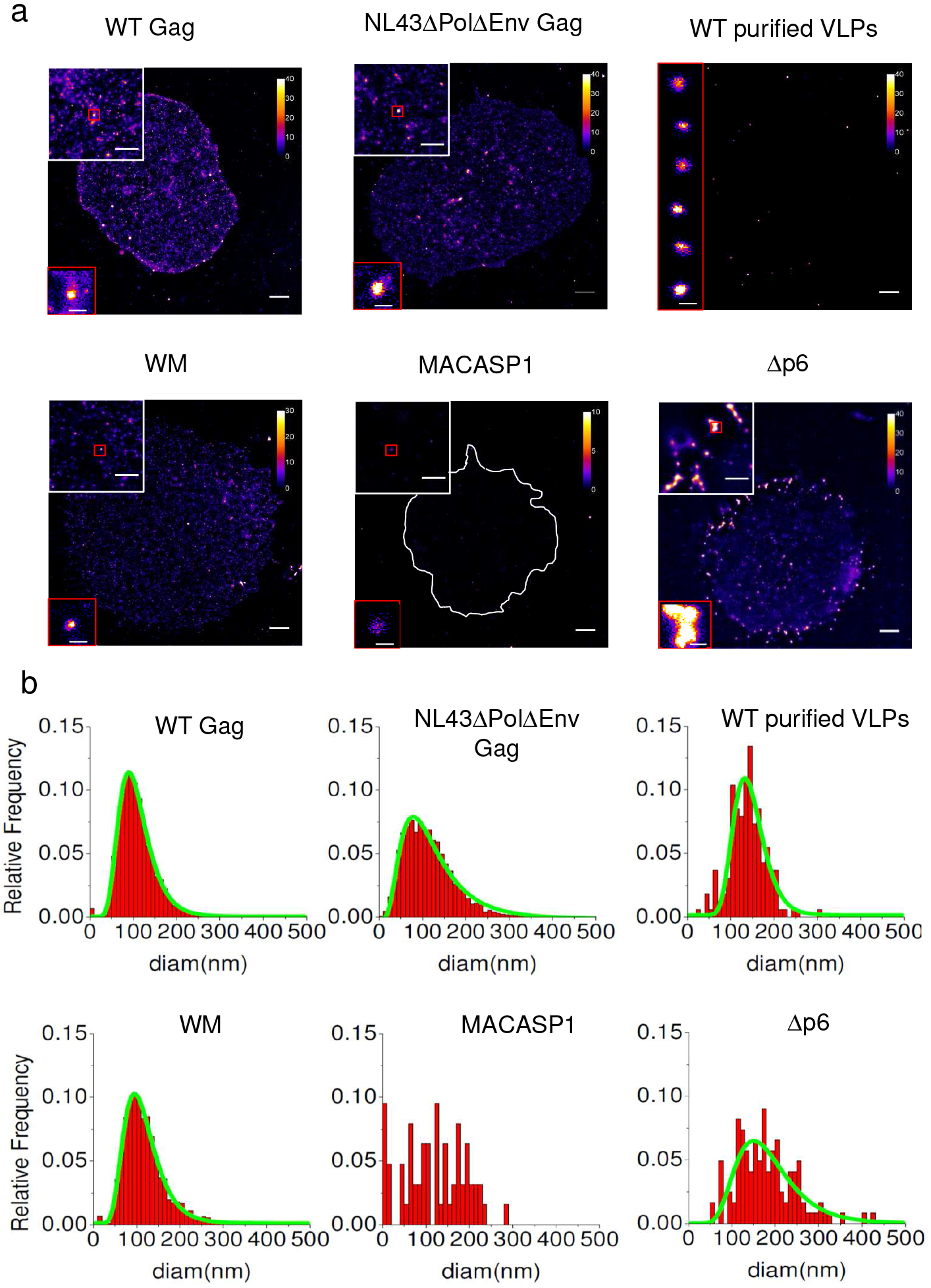
Global quantitative analysis of Gag(i)mEOS2 self-assembly at the surface of Jurkat T cells and in VLPs. (a) Jurkat T cells that express WT Gag, WM, MACASP1, Δp6 or NL4.3ΔPolΔEnv Gag were fixed and PALM imaging of the cell surface was performed using TIRF mode. Purified VLPs (left lower panel) from WT Gag productive cells were also analysed with this method and served as VLP size reference. Each image contains successive zooms of a selected area of the depicted cell (2x, upper left, 10x, lower left inset). Scale bars: 2, 1 and 0.2 *μ*m, respectively, from the largest field of view to the final zoom. (b) Size distribution (red) and their log-normal fit (green, when possible) of the different assembly clusters observed in T-cells. Value obtained from the fit are summarized in table 1.

**Table 1.** Mean Diameter and surface density of the identified clusters in Jurkat T cells: Quantification of Gag assembly platform size and density at the cell surface of fixed Jurkat T lymphocytes for WT Gag and each mutant (WM, MACA, Δp6), NL4.3ΔPolΔEnv Gag (in the presence of the packageable Psi-RRE genomic RNA), and the mean apparent VLP diameter. *: As illustrated in Figure S1, the Δp6 mutant is generating clusters of VLPs or arrested budding particles, which are not possible to resolve individually in PALM microscopy. Consequently, the mean density measured here is underestimated, and the mean apparent diameter is overestimated.

| | #clusters | Apparent Diameter (nm) (mean±sd) | n | Density ($\mu m^{-2}$) (mean±sd) |
| --- | --- | --- | --- | --- |
| WT Gag | 5089 | 112±54 | 5 | 4.5±1.7 |
| WM | 2627 | 121±56 | 5 | 1.7±1.5 |
| MACA | 63 | n.d. | 7 | 0.05±0.01 |
| $\Delta p6$ | 812 | 180±68* | 3 | 1.3±0.5* |
| NL43 ΔPolΔEnv Gag | 5239 | 117±65 | 4 | 3±1.2 |
| Purified VLPs | 164 | 139±42 | - | - |

### Quantifying the trapping energy experienced by single Gag molecules during VLP formation in living CD4 Jurkat T Cells

Since global analysis of Gag dynamics only partly reflected the properties of assembly, Live PALM was then used to monitor the changes in motion and densities of single Gag molecules in the vicinity of the assembling platforms. For that purpose, we created a movie with all the frames acquired in 26min. Each image of this movie included all the molecules localized in a 4min window with a sliding time of 10s. Figure 4a upper part shows two successive images of a such made movie of one forming VLP (WT Gag) exhibiting changes in molecular localization densities (LD) (see also figure S3). Figure 4a lower part show the associated trajectories of single WT Gag molecules around this forming VLP (see also video S1). In addition to diffusive motions, Gag trajectories directed towards the VLP centre were also observed, suggesting that VLPs in formation act as an energy trap for neighbouring Gag proteins. Therefore, Gag dynamics were analysed with the overdamped Langevin equation (suitable for heterogeneous diffusion processes) as an approximate model (Methods, eq.). A newly developed approach using Bayesian inference was used to analyse our data (23). Space was partitioned using a simple unsupervised approach and space domains were defined with the Voronoi tesselation. The estimators were the Maximum a Posteriori of the diffusion (D) and the effective trapping energy (E). Diffusion (Fig. 4b) and effective trapping energy (Fig. 4c) maps were built from the motions of all tracked single Gag molecules. To monitor precisely VLP formation, the start time and the position of forming VLPs (assembling platforms, hereafter) were identified as the time and position where the LD was three times higher than in the surrounding area. Moreover, the apparent VLP radius was also measured and was defined as the distance were the LD was four times lower than at the VLP centre (see Methods). By multiplexing this approach, more than 600 assembling platforms could be analysed in four different cells for WT and mutant Gag proteins. Then, LD, D and E changes during the overall acquisition time were investigated (see video S2 for examples of temporal changes of D and E maps) and were plotted (Fig. 4 d,e,f) for each forming VLP. From these plots, the maximal LD increase (LDI, red arrow) and the duration of this increase (dotted red arrow) could be measured. This last value was considered to represent the VLP formation time. It was previously shown (17) that HIV-1 Gag assembly in adherent HeLa cells can be divided in three phases characterized, respectively, by an increase in LD, followed by a plateau value and then a decrease in LD (due to particle release). These three phases could also be observed here in the host CD4 T cells (Fig. 4d, Table S2). The mean diffusion (bold dotted line in Fig. 4e, Methods, eq. 9) and the maximum trapping energy (ΔE, full red arrow in Fig. 4f, Methods, eq. 10) could also be extracted, as well as the duration of the trapping energy increase (dotted red arrow in Fig. 4f). As Gag self-assembly seemed to locally modify the mobility behaviour of the surrounding Gag molecules, first, diagrams of the normalized distributions of the ΔE and D values for WT Gag and mutants as well as for CAAX(i)mEos2 (negative control for assembly) were generated for all 600 assembly platforms identified in section 3 (Fig. 4g). These diagrams showed that the normalized distribution peaks were progressively drifting from the right for CAAX(i)mEos2 (non-assembling molecules) to the left for NL4.3ΔPolΔEnv Gag (assembling Gag molecules), i.e., from high to low diffusivity. Moreover, WM molecules formed two populations characterized by a distribution similar to that of MACASP1 and WT molecules, respectively. Importantly, while the Gag mutants had spread distributions, NL4.3ΔPolΔEnv Gag always exhibited a quite narrow and fairly centred peak. In parallel, the D/LDI diagram show that increasing LDI lead to decreasing D, supporting the usefulness of these 3 parameters to quantitatively describe the assembly process. Surprisingly, the mean ΔE values (1-2 kbT independently of the condition) (Fig. 4g) were not in favour of an attractive process during assembly. However, they were the mean of all “on-going” assembling VLPs, including those that will assemble imperfectly, i.e. not reaching the fully assembled VLP state. Indeed, the total density (i.e., LD sum over the acquisition time) should reach the maximum when VLPs are fully assembled, whereas the maximal LDI could be explained by different situations (see figure 5a for a schematic representation). For example, an assembly platform not leading to VLP formation can have a high LDI value, but a low total density (e.g., MACASP1 in Fig. 5b). Conversely, a VLP almost fully assembled suddenly appearing in the field of view will have a low LDI, but a high total density. To discriminate amongst these different possibilities, the LDI values were distributed as a function of the total density of every identified assembling platform (n >600) for each mutant (Fig. 5b). Finally, to monitor correctly the formation of single VLPs in T cells, only isolated assembling platforms (i.e., separated by 400 nm, which is about seven times the radius of a released VLP) were considered. We therefore explored threshold values for each parameter (LDI and total density value) in order to be in line with the relative assembling platforms densities observed in fixed cells (Fig. 2, table 1) and the VLP production (Fig. 1c). Particles with high LDI (>2,500 *μ*m^−2^) and total density values >20,000 *μ*m^−2^ were selected (red lines on fig 5b) since only two of the 600 MACASP1 clusters identified were above these threshold values. Conversely, these thresholds allowed selecting 91 assembly platforms for NL4.3ΔPolΔEnv Gag, 76 for WT Gag and 36 for WM. The reduction by half of the number of assembly platforms for WM compared with WT Gag is in agreement with their VLP release data (Fig. 1c) and cellular densities (Table 1). First, we analysed the distribution of the VLP formation time values (Fig. 5c) The results indicated that the mean time was not significantly different for WT Gag, NL4.3ΔPolΔEnv Gag and WM (1160±350s, 1020±370s and 1003±350s, respectively). However, the individual values were very variable (from 350s to 1800s). Calculation of the duration of the first two phases (i.e., LD increase and plateau) showed that the first phase (assembly) lasted about 5min for NL43ΔEnvΔPol Gag, 7 min for WT Gag and 6min for WM. The plateau phase duration was about 6 min for all three Gag variants (the exact values are in Supplementary Table 1). Altogether these results suggest that: i) MACASP1 forms low-density assembling platforms that mainly do not reach the VLP formation stage, ii) only WM and WT Gag can form high-density assembly platforms that lead to VLP formation, iii) on average, the time needed to make a VLP seems not affected by the presence of a packageable viral RNA or by CA-CA interactions.

**Fig. 4.**
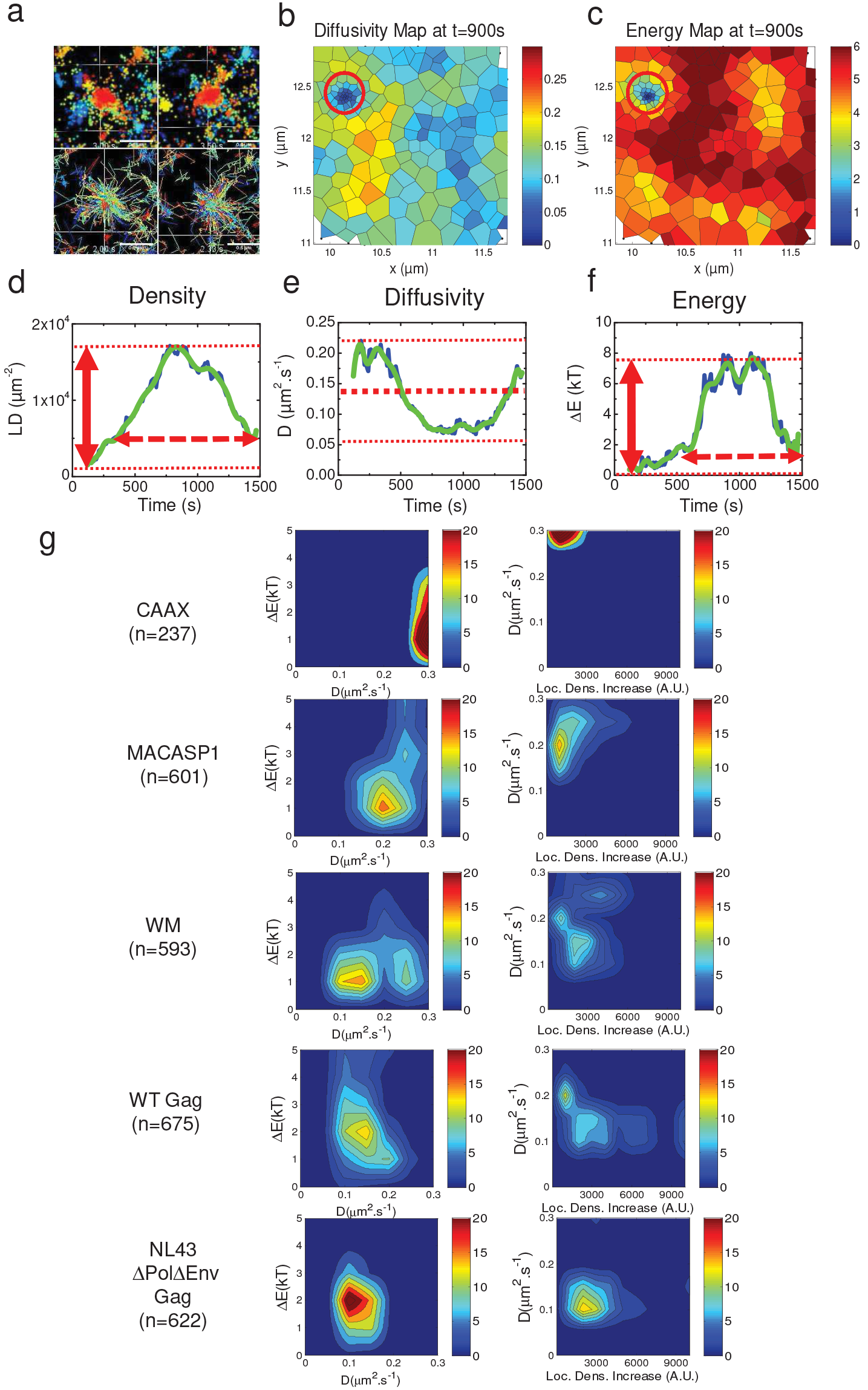
Monitoring time evolution of localisation densities, diffusivities and effective energies to characterize HIV-1 Gag(i)mEOS2 assembling VLP properties. (a) Typical examples of localisation changes overtime during assembly of WT Gag and as an illustration typical trajectories observed around an assembling VLP. All trajectories seem to be directed towards the centre of the assembling VLP. Every image represents all the localizations and trajectories observed in 4 minutes, (b, c) Diffusion (b) and effective trapping energy (c) maps obtained from multiple WT Gag dynamics analysis in a region of interest of the T cell membrane at t=900s of the experiment. Using Voronoi tessellation, the map was divided in sub-areas containing the same number of localizations (n=40). An assembling VLP is visible (inside the red circle) and can be identified by the increasing LD. (d) Typical plot of localization density (LD, *μ*m^−2^) variations overtime during assembly of one VLP. From this plot, the localization density increase (LDI, red arrow) and the assembly time length (red dotted arrow) were quantified for each assembling VLP for WT Gag and mutants. (e,f) The time-evolution of diffu-sion (e) and trapping energy (f) can also be plotted from the maps over the time length of the entire ex-periment to determine the mean diffusion constant (bold dotted line), the maximum trapping energy (red arrow) and the time length of trapping energy increase (dotted red arrow), x and y correspond to the position in *μ*m inside the PALM image, (e) Normalized distribution diagrams of the mean diffusion and the trapping energy (left), the mean diffusion and the localisation density increase (right) for each experimental condition, from CAAX(i)mEOS2 (no assembly) to WT Gag and NL4.3ΔPolΔEnv Gag (highest assembly efficiency).

Then, we quantified the effective trapping energy for the VLP releasing assembly platform (Fig. 5d). The only two selected MACASP1 particles exhibited no difference in effective trapping energy values compared with the total pool shown in Fig. 5d (ΔE=1.5 ± 0.1 *k*_*b*_*T*), whereas, the maximal effective trapping energy value slightly increased for the subset of VLP generated by WM (ΔE=2.1 ± 1.2 *k*_*b*_*T*) compared with the total pool. Conversely, the effective trapping energy value was strongly increased for WT Gag (ΔE = 3.7 ± 1.6 *k*_*b*_*T*) and NL4.3ΔPolΔEnv (ΔE = 3.7 ± 2.1 *k*_*b*_*T*) compared with WM (p<5.10^−5^, Student’s t test). These data indicate that a lack of correct CA-CA interactions during assembly induced a decrease of 40% in effective Gag trapping energy, ie Gag interaction, whereas the loss of NC abolished the existence of a trapping energy. Interestingly, the presence of the packageable viral RNA, or a Gag optimized codon sequence, did not influence the mean effective trapping energy value questioning the viral RNA assembly catalyser role. Indeed, molecular self-assemblies are more efficient when triggered by heterogeneous seeds (31–33), ie. Gag(NC)-RNA interaction could here act as the necessary seeds for controlling Gag self-assembly. To test this hypothesis, the time to reach the maximum LDI was compared to the time to reach the maximal effective Gag trapping energy intensity during the VLP assembly, for WM, WT Gag and NL4.3ΔPolΔEnv Gag. Interestingly, differences in time were dispersed both for WM and WT Gag, whereas NL4.3ΔPolΔEnv Gag showed one major peak centred on 0 (Fig. 5e). This important result shows that most probably the presence of the packageable *Psi*-containing viral RNA genome, containing a wild-type Gag Rev-dependent coding sequence, acts as a seed that favour the temporal and spatial coordination of viral Gag particle assembly at the host CD4 T cell plasma membrane.

## Discussion

In the last ten years, many efforts have been made to study the assembly of HIV-1 Gag particles in living cells (16–19, 34). Here, we monitored immature HIV-1 assembly dynamics in the host CD4 T lymphocytes using quantitative live PALM imaging and advanced big data quantitative analyses, , and we thus highlighted the role of CA protein-CA protein and NC protein-RNA interactions using different known Gag assembly defective mutants. Analysis of the effects of different Gag mutations on viral assembly platform formation at the cell surface of Jurkat T cells showed that the assembly platform size was on average bigger for CA(WM) mutant compared to WT Gag. Assembly platform densities were also quite different among the Gag proteins tested, reflecting their efficacy in binding to or assembling at the cell membrane (Fig.S2 and 2). MACASP1 molecules showed the most drastic assembly platform density reduction (by 50-fold compared with WT Gag) while WM exhibited only a 3-fold reduction. This suggests that the assembly efficiency at the T cell membrane strongly depends on Gag C-terminal domains (NC-sp2-p6), but not on the CA-SP1 interface alone. This is in good agreement with the work of Robinson *et al.* (35), showing that a MACA mutant is unable to produce high order multimers of Gag using velocity sedimentation and gradient assays. Furthermore, the 20-fold higher density for Δp6 compared with MACASP1 suggests that a direct effect of ESCRT protein (such as Tsg101-p6 interaction) on assembly platform formation is unlikely. In addition, VLP formation analysis (videos S3 and S2 for Δp6 and WT Gag, respectively) showed that Δp6 Gag molecules can assemble, but accumulate in the same location. Δp6 Gag signal persistence might indicate a defect in particle release, compared with WT Gag (as it can be visualized by electron microscopy in (14) and confirmed here for Δp6 Gag(i)mEOS2 (see Fig S1)). We thus propose that the remaining NC domain is the main determinant for Gag assembly platform formation at the surface of Jurkat T cells, most probably by a trapping of the Gag molecules via NC-RNA interactions that induce Gag-Gag multimerization. To go further, we performed live PALM and quantify single-molecule dynamics by analysing the trajectories of individual Gag molecules. We first characterized the motion of individual molecules using a simple Brownian diffusion approach. By analysing tens of thousands trajectories, extracting and decomposing their diffusion coefficients distribution, we detected that the proportion of assembling molecules was the highest for WT Gag and NL4.3ΔPolΔEnv Gag in Jurkat T cells. Conversely, for oligomerization-deficient mutants (WM and particularly MACASP1), the proportion of highly mobile, non-assembling molecules was the highest. This confirms that, these two mutants are less engaged in platform formation at the surface of Jurkat T cell plasma membrane. Hendrix *et al.* also observed an increase of the mobile fraction of oligomerization-deficient mutants in the cytosol (29). Although we performed live PALM with TIRF illumination, we cannot exclude that a cytosolic fraction of Gag molecules linked to RNA might contribute to the measured diffusion coefficients. Nevertheless, in this case, we would also expect to observe the fast diffusing cytosolic component of RNA-attached WT Gag (D=2.8±0.5 *μm*^2^.*s*^−1^) described by Hendrix *et al.* (29). Our D distributions show that this component is poorly present, suggesting that the observed Gag oligomerizing fraction was mainly located at the plasma membrane of Jurkat T cells. Then, we used quantitative live PALM on Gag(i)mEOS2 proteins in Jurkat T-cells to monitor their spatial density increase over time and to decipher the kinetics of viral particle formation. We observed three different phases in the formation and release of newly formed particles, as previously described in adherent HeLa cancer cell lines by TIRF microscopy. By live PALM in TIRF configuration, we found for WT Gag and NL4.3ΔPolΔEnv Gag, that the first phase (considered as the assembly phase) lasted approximately 5min, in agreement with the value found by Jouvenet *et al.* (16) but lower than what reported by Ivanchenko *et al.* (17) (both in adherent HeLa cells). The second phase (a plateau phase, supposedly when the ESCRT machinery is recruited (36)) lasted approximately 10min, as observed in Hela cells (17). We did not quantify the third phase because it was quite variable, possibly due to many different processes (pinching off, bleaching…), but instead measured the total time of particle formation as the time length between the appearance and disappearance (i.e., return to the initial LD) of an assembly platform (see video S1). The mean duration was observed between 17min (for NL4.3ΔPolΔEnv) and 20min (for WT Gag), but with high variability (7 to 30min), as already observed in HeLa cells (36). This shows that the presence of the viral RNA does not significantly decrease the total assembly and release time of the virus like particle. Moreover, our and previous results (17, 36), show that assembly duration and the time needed to achieve a fully released particle seem to be independent of the cell type. While visualizing individual Gag molecule tracks, we could observe that WT Gag(i)mEOS2 proteins tended to move towards the centre of assembling particles, suggesting the existence of an attractive potential. However, this could be an artefact due to the high molecule density in the platforms, leading to systematic reconnections with the molecules moving in their proximity (37). As full trajectories were not required for VLP mapping, image to image graph matching was used to generate the most probable protein displacements (38). The live PALM data were then analysed using Bayesian inference and the modified Langevin equation to quantify the motion of individual Gag molecules (23, 24, 39). Using the Langevin description of the motion, the key dynamical properties were approximate as diffusion and effective energy maps, providing us with a more general understanding of the modifications of protein dynamics at the vicinity of assembling platforms. Therefore, by computing temporal maps of the diffusion and energy properties in and around the assembling platforms, we could measure the intensity and the spatial range of this attracting energy. We obtained the highest intensity for NL4.3ΔPolΔEnv Gag and WT Gag (< Δ*E* >~ 4*k*_*b*_*T*). This value was almost half for the CA-CA(WM) mutant molecules that still generate platforms and was down to 1.5 *k*_*b*_*T* for MACASP1 molecules that do not generate VLPs. This last value is close to the thermal fluctuation energy (1 *k*_*b*_*T*) and therefore, typical of an energy that induced no Gag trapping. Recently, using coarse grained molecular simulation Pak *et al.* (40) has estimated that the oligomerization via CA-CA hexamer formation due to SP1 interactions in the presence of (simulated) membrane and RNA could occur for weak SP1-SP1 interaction energy (~4 *k*_*b*_*T*). Interestingly, this value is exactly in line to what we find in Jurkat T-cells in the case of WT Gag. Nevertheless, the SP1-SP1 interaction defective mutant (WM) still generates virus like particules in CD4 T-cells although exhibiting a < ΔE > value lower than 4*k*_*b*_*T*. Moreover, the long spatial range (200nm, i.e., 1.5 bud diameter) observed here of the attractive forces apparently generated by assembly, suggest that this mean effective Gag trapping energy field cannot only be due to direct protein/protein or protein/RNA molecular interactions. On the opposite, membrane curvature-mediated interactions and forces exerted by each protein on the cell membrane could account for the energy fields observed here. Sens et *al.* (41,42) theoretically predicted for caveolae that, when the force exerted on the plasma membrane by oligomers decreases, the resulting bud radius is proportionally increased. In our study, the almost two-fold reduced attractive energy for WM (reduced SP1-SP1 interactions) was correlated with the almost two-fold increase in the VLP diameter observed by electronic microscopy (Fig. S1). Results obtained here suggested that in Jurkat T-cells, the plasma membrane can then rescue the lack in efficient CA-CA dimerization in order to produce immature particles, as already observed on model membranes (9). This is also in good agreement with the labile membrane bound form of WM Gag mutant observed in 293T HEK cells in the Robinson *et al.* (35) biochemical and EM study. Finally, nor the presence of the assembly-triggering Psi RNA sequence, neither the Rev/RRE driven RNA trafficking or codon optimized Gag sequence, did change the attraction energy intensity. Therefore, to gain more insights into the role of CMV-driven codon optimized Gag (WT Gag and WM) versus the cis packageable Rev/RRE driven viral Psi containing RNA genome (NL4.3ΔPolΔEnv Gag), we analysed the temporal correlation between Gag density and the Gag attraction energy increase. Unlike WT Gag and WM, a perfect temporal correlation between Gag density and Gag trapping energy was observed for NL4.3ΔPolΔEnv Gag. This result reveals an important role for the Psi/RRE containing viral RNA genome in the spatio-temporal coordination of HIV-1 Gag assembly at the plasma membrane of CD4 T cells. It could be due either by contributing to the specific interaction of the genomic RNA with the NC domain of Gag during Gag multimerization at the cell membrane, or by the fact that a Rev/RRE genomic RNA intracellular trafficking and location will contribute to coordinate virion assembly. This latter hypothesis goes with the recent work of Becker and Sherer showing that the viral mRNA subcellular trafficking and location regulates viral assembly at the cell membrane (19). The first hypothesis is in good agreement with the previous finding (43, 44) that the viral RNA genome (containing the Psi signal for encapsidation) acts as a structural element for retroviral particles, and with the model recently proposed by Chen *et al.* (26) showing that miRNA binding to Gag NC inhibits HIV-1 assembly. This goes with reports showing that high-order Gag multimerization only occurs at the cell membrane (45) and is dependent on Gag membrane-binding capacity. Conversely, low-order Gag multimers bind to viral genomic RNA in the cytosol prior to assembly and would be dependent on the NC domain of Gag (29, 35, 45). Here, our results suggest that the formation of high-order Gag multi-mers occurring at the plasma membrane of CD4 T cells depends strongly on NC.

**Fig. 5.**
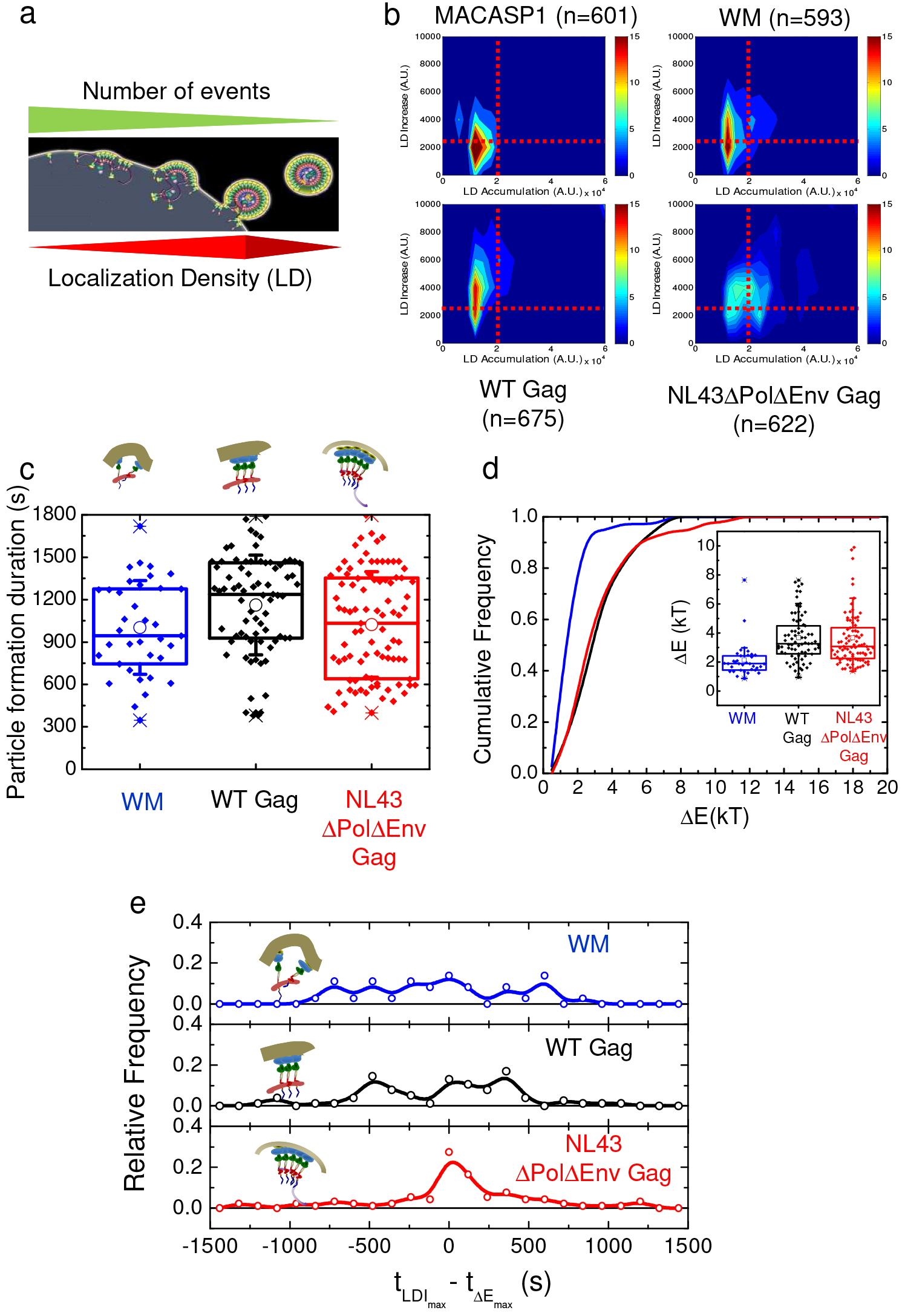
Maximal effective trapping energy and diffusion of Gag(i)mEOS2 (WT and mutants) in VLPs during their assembly at the plasma membrane of Jurkat T cells. (a) Schematic representation of VLP assembly showing the different steps and their relation with LD changes. The final step (before VLP release) is expected to be the densest step (LDI) and the less frequent one. (b) Diagram of LDI as a function of total LD (LD accumulation during the entire experiment) for WT Gag, the WM and MACASP1 mutants, and NL4.3ΔPolΔEnv Gag. n: number of VLPs before selection of assembling VLPs; red lines delimit the threshold value used for both LDI and total LD to sort the fully assembling VLPs from the others. (c) Gag(i)mEOS2 assembly duration distribution for WM, WT Gag and NL4.3ΔPolΔEnv Gag after thresholding. The mean value of particle formation duration was 16±6min for WM (n=36), 20±6min for WT Gag (n=76) and 17±6min for NL4.3ΔPolΔEnv Gag (n=91). (d) Cumulative frequency of the maximum effective trapping energy obtained with fully assembling VLPs. The cumulative frequency was obtained from the distribution shown in the inserted box-plot. (e) Normalized distributions of the difference between the time to reach the maximum LD and the time to reach the maximal energy trapping for WM (blue), WT Gag (black) and NL4.3ΔPolΔEnv Gag (red).

**Fig. 6.**
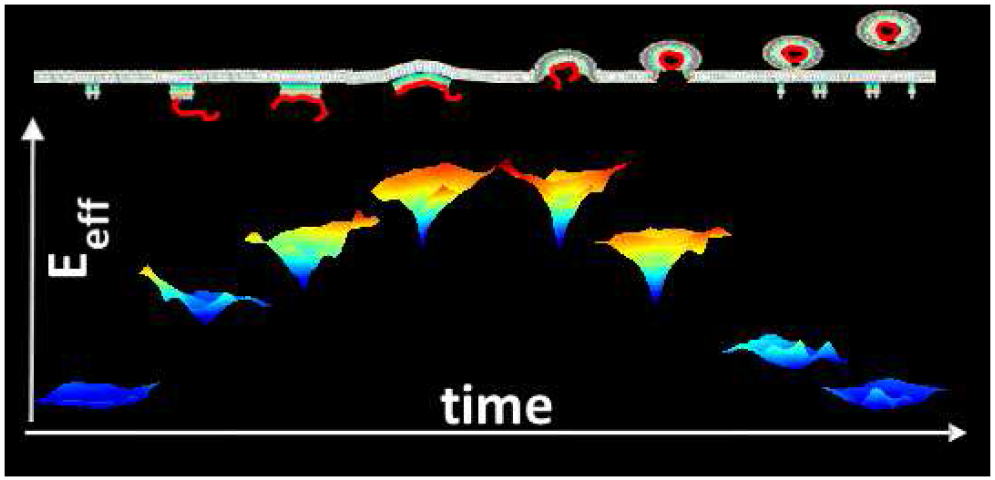
A schematic view of change in effective Gag trapping energy during HIV Gag assembly and budding. The top scheme represents Gag molecules oligomerization at the cell plasma membrane (in grey) with the viral RNA genome (in red). Below is represented the color coded (increasing from blue to red) effective attractive energy well. Deepest depth represent the most attractive situation.

In conclusion, using quantitative single molecule microscopy, we could measure the evolution of the trapping energy occuring during Gag self-assembly at the CD4 T cell plasma membrane (see figure 6 for a schematic illustration). We were able to show, for the first time, that the presence of a packageable cis-acting viral RNA genome, coding for Gag, revealed a spatio-temporal coordination of HIV-1 assembly at the CD4 T cell plasma membrane. This strongly suggests that the faith of the viral RNA genome (trafficking or encapsidation) is synchronizing Gag assembly at the cell membrane.

## Methods

### Cell culture

Jurkat T cells (a human T-cell leukaemia cell line) were grown in RPMI 1640 plus Glutamax (Gibco) supplemented with 10% foetal calf serum (FCS) and antibiotics (penicillin-streptomycin. Human embryonic kidney (HEK) 293T cells were grown in DMEM (Gibco) supplemented with 10% FCS and antibiotics (penicillin-streptomycin).

### DNA plasmids

The plasmid expressing HIV-1 Gag with the internal (located between MA and CA) mEOS2 tag fused with the Gag protein was called pGag(i)mEOS2 WT (CMV promoter-driven codon optimized Gag sequence), and used to generate the WM and MACASP1 mutants by site-directed mutagenesis. Another plasmid that expresses HIV-1 WT Gag(i)mEOS2 and the viral cis-packageable Psi RNA genome with a Rev/RRE dependent Gag coding sequence (pNL4.3ΔPolΔEnv) was kindly provided by Dr Eric Freed (NIH, Frederick, MD, USA), and was described previously (26).

### Site-directed mutagenesis

Mutations were introduced in pGag(i)mEOS2 WT by site-directed mutagenesis using the QuickChange mutagenesis kit (Agilent) according to the manufacturer’s protocol. Tryptophan 184 and methionine 185 were replaced by two alanine residues (WM mutant, as in (29)) using the primer 5’-GAC GTG AAG AAC GCA GCT ACC GAG ACC CTG-3’. NC-sp2-p6 were deleted by inserting a stop codon after the sp1 sequence (MACASP1 mutant) using the primer 5’-GCG ACC ATC ATG TAG CAG CGC GGC AAC-3’. The p6 sequence was deleted by inserting a stop codon after the sp2 sequence (Δp6 mutant) using the primer 5’- CCC GGC AAC TTC TAG CAG AGC CGC CCC-3’. All plasmids were amplified in E. coli and mutations were confirmed by DNA sequencing (MWG Eurofins).

### DNA transfection

Jurkat T cells (2×106) were microporated with 4μg of each plasmid using the Amaxa system (Lonza) and then plated in RMPI complete medium and harvested 24h post-transfection, as described in (27). HEK293T cells were transfected by using the calcium phosphate buffer, as described in (46).

### Cell viability, transfection efficiency and protein expression by FACS

Cell viability, cell transfection and protein expression were assessed with a BD FACS Calibur flow cytometer. FACS results were analysed with the FlowJo software v10. The cell viability rate was calculated as the ratio of cell size and granulometry over the total cell number. Protein expression rates were monitored using the geometric mean of the mEOS2(+) cell fluorescence intensity distribution. Antibodies. Western blots were performed using the anti-CAp24 (NIH AIDS Reagent Program HIV-1 p24Gag monoclonal (24-4) mouse antisera) and mouse anti-LAMP2 (human lysosome-associated membrane protein 2) (H4B4) (Santa Cruz Biotechnologies) antibodies, followed by anti-mouse and anti-rabbit antibodies coupled to horseradish peroxidase (HRP) (Dako), and an anti-GAPDH HRP-coupled antibody (Abcam).

### VLP purification and immunoblotting

To monitor viral particle production, cell culture media containing VLPs were harvested at 24h posttransfection. Jurkat T cell supernatant was clarified by centrifugation at 60xg for 10min, while HEK293T cell supernatants were filtered (0.45 μm pore size). Viral supernatants were purified by ultracentrifugation through a sucrose cushion [25% glucose (wt/vol) in TNE buffer (10mM Tris-HCl pH 7.4, 100mM NaCl, 1mM EDTA)], at 100000xg in a Beckman MLA150 or SW60Ti rotor for 90min. ellets were resuspended at 4°C in TNE buffer overnight and stored at −80°C. To analyse the intracellular viral protein content, cells were lysed in RIPA buffer —(150mM NaCl, 20mM Tris-HCl [pH 8], 1% NP-40, 0.1% SDS, 0.2mM EDTA) and sonicated. Cell lysates were then clarified at 16000xg for 10min, and the protein concentration in the cell lysates was determined with the Bradford assay. For western blot analysis, 50 μg of total proteins were loaded and separated on 10% SDS-PAGE gels and transferred onto polyvinylidene difluoride membranes (Thermo Fischer). Immunoblotting was performed using the relevant antibodies and HRP signals were revealed with the SuperSignal West Pico substrate (Thermo Scientific). Transfection efficiency and VLP release calculation. Plasmid transfection efficiency in T cells was evaluated by measuring the percentage of fluorescent cells by immunofluorescence or flow cytometry analysis. For VLP release, the HRP signals from immunoblot membranes were imaged using the G:Box system (Syngene), and the viral Gag or CA protein signals were quantified using the ImageJ software. The percentage of VLP release relative to the GAPDH loading sample was then estimated as:

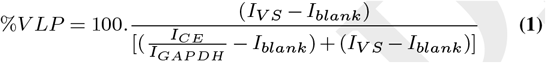

where *I*_*V S*_ is the viral supernatant (quantification of the immunoblot signals for Gagp55 and CAp24); *I*_*C E*_ is the quantification of the Gagp55 signal in cell extracts; and *I*_*blank*_ is the background signal. GAPDH served as loading control.

### Photo Activation Localization Microscopy

#### Sample Preparation

To increase cell adhesion, Jurkat T cells were seeded on poly-lysine-coated coverslips and left in culture medium at 37°C, 5% CO2 for 30 min. After rinsing with PBS, cells were incubated at 37°C in microscopy buffer (MB) (150mM NaCl, 20mM HEPES pH7.4,1mM CaCl2, 5mM KCl, 1mM MgCl2 pH 7.4 and 100 nm TetraSpeck™ microspheres) for live PALM experiments, or fixed with 3% PFA in PBS at room temperature for 15min for PALM experiments. In this case, after fixation, cells were rinsed with 50mM NH4Cl for 5min and then several times with PBS before transfer in MB for microscopy analysis.

#### Live PALM and PALM image acquisition

Cells were imaged at 37°C in Ludin chambers (Life Imaging Services) with an inverted motorized microscope (Nikon Ti) equipped with a 100x 1.45 NA PL-APO objective and a perfect focus system for long acquisition under TIRF illumination. For live PALM, cells that express mEOS2-tagged Gag molecules were photoactivated using a 405nm laser (Omicron) and the resulting photoconverted single-molecule fluorescence was excited with a 561nm laser (Cobolt Jive™). Both lasers illuminated the sample simultaneously. The photoactivation laser power was adjusted to keep the number of the stochastically activated molecules constant and sparsely distributed during the acquisition to allow single-molecule localization (46). Fluorescence signals were collected by using a dichroic and an emission filter (F38-561 and F39-617, respectively, Semrock) and a sensitive EMCCD camera (Evolve, Photometric). Acquisition was guided by the MetaMorph software (Molecular Devices) at 50Hz in streaming mode and analysed online with laser feedback to ensure the optimal and constant number of localizations during acquisition (46). Multicolour fluorescent 100nm TetraSpeck™ microspheres (Invitro-gen) were used as fiduciary markers to acquire and correct for lateral drifts occurring during long-term acquisitions.

#### Single-molecule acquisition and tracking

LivePALM experiments allowed the acquisition of sets of 80,000 images per cell that were analysed with WaveTracer (Molecular Devices) and the custom-made PALM Tracer analysis software to extract molecule localization and dynamics data. Fluorescent single molecules were localized and tracked over time using a combination of wavelet segmentation and simulated annealing algorithms (47), (48) operating as a plug-in for the MetaMorph software (Molecular Devices). Using the same experimental conditions described above, the system resolution was quantified to 46 nm at full width and half maximum using fixed mEOS2-expressing cells. 200 two-dimensional distributions of singlemolecule positions belonging to long trajectories (>30 frames) were analysed by bi-dimensional Gaussian fitting. The resolution was defined as 2.3 *σ*_*xy*_, where *σ*_*xy*_ was the standard deviation of the Gaussian fit. Trajectories equal or longer than 10 points were analysed using the mean squared displacement MSD computed as

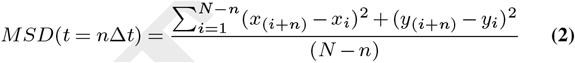

where *x*_*i*_ and *y*_*i*_ were the coordinates of the label position at the time *iΔt.* The diffusion coefficient D was defined as the slope of the affine regression line fitted from the MSD(nΔ t) with 0 < *n* < 5. The log distribution of D experimental values was linearly decomposed by the log distribution of VLP and CAAX(i)mEOS2 using the following equation:

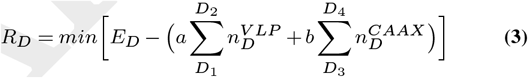

where *R*_*D*_ was the residual, *E*_*D*_ the experimental distribution, and *a* and *b* the proportions of VLP and CAAX distributions in each experimental distribution. *D*_1_, *D*_2_ *D*_3_ and *D*_4_ were respectively set to: 10^−5^, mean VLP value (2.10^−3^), mean CAAX value (0.6), 20 *μm*^2^.*s*^−1^. Assembling platform apparent diameter were determined as the 1/*e*^2^ diameter of a bi-dimensional Gaussian fitting.

#### Robust statistical Gag dynamics analysis using Bayesian inference

The large amount of data and the time-evolving nature of the process required an automated and stereotypical way to treat data. Thus, all data were treated exactly in the same manner, including the inference hyperparameters. A pipeline was designed for all data analysis. The pipeline included five steps:

- Single Molecule Localization

- Non-tracking with Graph Assignment

- Selection of the Regions of Interest

- Time-Evolving Bayesian Inference Analysis

- Time-Evolving Feature Extraction from Inferred Maps

Single Molecule Localization was implemented in MATLAB using the slightly modified MTT algorithm (49). To limit errors due to tracking algorithms, we did not track single molecules, but used optimal assignments between consecutive images to extract Gag movements. This procedure is detailed in Supplementary Note **??**. Region of interest (ROI) selection was then based on LDs (localisation densities). ROIs were selected as squared areas of 2 *μm* in length centred on the maximum of density. The number of ROIs per cell was limited to 30. In each ROI, the effective centre of a VLP, *r*_*eff*_, was defined as the point with the highest LD 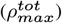 (cumulated on the 80 000 frames). The effective radius, *R*_*eff*_, of a VLP was defined as the average distance between *r*_*eff*_ and the points of a density equal to 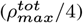. In the analysis, points within *R*_*eff*_ were considered to be in the VLP. Depending on the density of maturing VLPs, more than one VLP could be present in a single ROI. All VLPs inside such region were analysed. They were discarded afterwards by LD/LDI selection (figure 5). Then, the molecule motion was analysed using Bayesian inferences (see Supplementary Methods for details). Briefly, the dynamics of individual Gag proteins were approximated with the Overdamped Langevin Equation (OLE) written using an Itô interpretation:

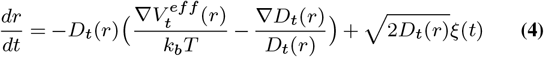

where *D*_*t*_(*r*) was the space-varying diffusivity, 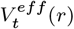 the effective potential, and *ξ*(*t*) the zero-averaged Gaussian noise. 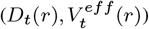 were
considered to be the statistical features that encode the dynamical characteristics of the environment concerning the individual dynamics of Gag proteins. Bayesian Inference was used to extract 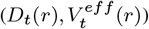 from the assignment between images (23, 50–53) (see Supplementary Methods for details of the Bayesian procedure). In equation, the index t shows that diffusion and potential fields can change with time, but at a time scale larger than that of the particle dynamics. Nevertheless, the time-evolving dynamics during VLP assembly led to high variability in particle density. Thus, the tessellation procedure described in (23) was modified to ensure more homogeneity in the structure of the Voronoi spatial tessellation (details in Supplementary Methods). Finally, temporal changes in VLP dynamics were monitored by time windowing. The duration of the window was set to 240s with a sliding time of 10s. The inference was performed independently for each time window, allowing the map to be inferred independently of previous or future detections. Hence, the typical time-evolution of VLP maturation led to 136 maps of 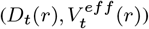. VLP density inside a map (corresponding to a time window) was evaluated as

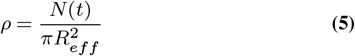
where *N*(*t*) was the number of localizations in the VLP during the time window t. The time-evolution of density for a VLP was directly computed as the density measured on the set of maps associated with that VLP. Considering that the set of mesh subdomains in the VLP at the time window t is *I*(*t*) and neighbours to a VLP are defined as the set of mesh subdomains in contact with the VLP and referred as *M*(*t*), then, at each time window t, the diffusivity in the VLP was defined as

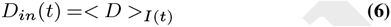
and the VLP trapping energy was defined as

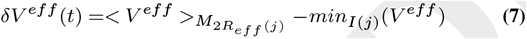
where <.> was the spatial average. Finally, for each VLP present in the analysed ROIs (i.e., 600 VLPs per mutant), the time-evolving density, diffusivity and potential were smoothed with a 10th order Savitsky-Golay filter to extract the following parameters:

- Localization Density Increase (LDI):

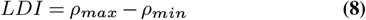

- Mean diffusivity in the assembly platform :

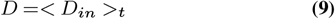

where <.> was the time average.

- Maximum Trapping Energy :

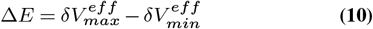

The assembly time length was defined as the 1/*e*^2^ width of the Gaussian fit of the LDI peak. As the analysis led to ≃ 100,000 maps of 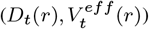, the 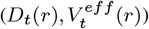 values observed in each VLP were distribute in classes. Each class was renormalized to the total VLP number. This allowed generating (*V*^*eff*^ = *f*(*D*),D = *f*(*LDI*)) diagrams for each mutant.

## ACKNOWLEDGEMENTS

We are thankful to Dr. Suliana Manley (EPFL, Switzerland) and Julia Guzen-hauser for helpful discus-sions. We are thankful to Dr. Eric Freed for the gift of the pNL43ΔPolΔEnv plasmid. J.B.S., J.B.M., M.D., C.F. and D.M. are part of the CNRS GDR ImaBio consortium. We also thank MRI and CEMIPAI platforms for the access to the PALM/TIRF microscopes in Montpellier (France). This work has been funded by ANR Fluobuds (grant ANR-13-BSV5-0006-01). C.FLo fellowships were funded by the GDR ImaBio for the Master and the University of Montpellier for the PhD. This work was also supported by the regional council of Aquitaine, the LabEx BRAIN and the Fondation pour la Recherche Médicaleto JBS. This workwas funded by the state program Investissement d’avenir managed by the ANR (grant ANR-10-BINF-05 “Pherotaxis”), the ANR TRamWAy and the Pasteur start package of the Decision and Bayesian Computation laboratory to JBM.

## AUTHOR CONTRIBUTIONS

DM designed the study. DM and CF wrote the manuscript with the help of JBS and JBM. CFlo, JBS and CF performed PALM and Live PALM experiments. SG and PR performed and analyzed EM experiments. CFlo, PM, EB and DM performed cell biology, biochemical analysis and plasmid design. CFlo, JBS and CF analyzed live-PALM and PALM data. JBM wrote the pipeline in C_++_ and Matlab to extract and analyze live PALM data, with participation of MD and MEB. CF wrote Matlab scripts for live PALM data interpretation and representation.

## Bibliography

1 Suliana Manley, Jennifer M. Gillette, George H. Patterson, Hari Shroff, Harald F. Hess, Eric Betzig, and Jennifer Lippincott-Schwartz. High-density mapping of single-molecule trajectories with photoactivated localization microscopy. Nature methods, 5:155–157, February 2008. 10.1038/nmeth.1176.

2 Eric Betzig, George H. Patterson, Rachid Sougrat, O. Wolf Lindwasser, Scott Olenych, Juan S. Bonifacino, Michael W. Davidson, Jennifer Lippincott-Schwartz, and Harald F. Hess. Imaging intracellular fluorescent proteins at nanometer resolution. Science (New York, N.Y.), 313(5793):1642–1645, September 2006. ISSN 1095-9203. 10.1126/science.1127344.

3 Eric O. Freed. HIV-1 assembly, release and maturation. Nature reviews. Microbiology, 13: 484–496, August 2015. 10.1038/nrmicro3490.

4 D. Gheysen, E. Jacobs, F. de Foresta, C. Thiriart, M. Francotte, D. Thines, and M. De Wilde. Assembly and release of HIV-1 precursor Pr55gag virus-like particles from recombinant baculovirus-infected insect cells. Cell, 59(1):103–112, October 1989. ISSN 0092-8674.

5 Marilyn D. Resh. Intracellular trafficking of HIV-1 Gag: how Gag interacts with cell membranes and makes viral particles. AIDS reviews, 7(2):84–91, June 2005. ISSN 1139-6121.

6 Vineela Chukkapalli and Akira Ono. Molecular determinants that regulate plasma membrane association of HIV-1 Gag. Journal of Molecular Biology, 410(4):512–524, July 2011. ISSN 1089-8638. 10.1016/j.jmb.2011.04.015.

7 Charlotte Mariani, Marion Desdouits, Cyril Favard, Philippe Benaroch, and Delphine M. Muriaux. Role of Gag and lipids during HIV-1 assembly in CD4(+) T cells and macrophages. FrontiersinMicrobiology, 5:312, 2014. ISSN 1664-302X. 10.3389/fmicb.2014.00312.

8 Siddhartha A. K. Datta, Joseph E. Curtis, William Ratcliff, Patrick K. Clark, Rachael M. Crist, Jacob Lebowitz, Susan Krueger, and Alan Rein. Conformation of the HIV-1 Gag protein in solution. J Mol Biol, 365(3):812–824, January 2007. 10.1016/j.jmb.2006.10.073.

9 Naresh Yandrapalli, Quentin Lubart, Hanumant S. Tanwar, Catherine Picart, Johnson Mak, Delphine Muriaux, and Cyril Favard. Self assembly of HIV-1 Gag protein on lipid membranes generates PI(4,5)P2/Cholesterol nanoclusters. Scientific Reports, 6:39332, December 2016. ISSN 2045-2322. 10.1038/srep39332.

10 Hanumant S. Tanwar, Keith K. Khoo, Megan Garvey, Lynne Waddington, Andrew Leis, Marcel Hijnen, Tony Velkov, Geoff J. Dumsday, William J. McKinstry, and Johnson Mak. The thermodynamics of Pr55gag-RNA interaction regulate the assembly of HIV. PLoS pathogens, 13:e1006221–e1006221, February 2017. 10.1371/journal.ppat.1006221.

11 Alan Rein, Siddhartha A. K. Datta, Christopher P. Jones, and Karin Musier-Forsyth. Diverse interactions of retroviral Gag proteins with RNAs. Trends in Biochemical Sciences, 36(7): 373–380, July 2011. ISSN 0968-0004. 10.1016/j.tibs.2011.04.001.

12 Delphine Muriaux and Jean-Luc Darlix. Properties and functions of the nucleocapsid protein in virus assembly. RNA biology, 7(6):744–753, December 2010. ISSN 1555-8584.

13 Hugues de Rocquigny, Salah Edin El Meshri, Ludovic Richert, Pascal Didier, Jean-Luc Darlix, and Yves Mély. Role of the nucleocapsid region in HIV-1 Gag assembly as investigated by quantitative fluorescence-based microscopy. Virus Research, 193:78–88, November 2014. ISSN 1872-7492. 10.1016/j.virusres.2014.06.009.

14 Uta K. von Schwedler, Melissa Stuchell, Barbara Müller, Diane M. Ward, Hyo-Young Chung, Eiji Morita, Hubert E. Wang, Thaylon Davis, Gong-Ping He, Daniel M. Cimbora, Anna Scott, Hans-Georg Kräusslich, Jerry Kaplan, Scott G. Morham, and Wesley I. Sundquist. The protein network of HIV budding. Cell, 114(6):701–713, September 2003. ISSN 0092-8674.

15 Siddhartha A. K. Datta, Lakew G. Temeselew, Rachael M. Crist, Ferri Soheilian, Anne Kamata, Jane Mirro, Demetria Harvin, Kunio Nagashima, Raul E. Cachau, and Alan Rein. On the role of the SP1 domain in HIV-1 particle assembly: a molecular switch? Journal of Virology, 85(9):4111–4121, May 2011. ISSN 1098-5514. doi: 10.1128/JVI.00006-11.

16 Nolwenn Jouvenet, Paul D. Bieniasz, and Sanford M. Simon. Imaging the biogenesis of individual HIV-1 virions in live cells. Nature, 454(7201):236–240, 2008.

17 Sergey Ivanchenko, William J. Godinez, Marko Lampe, Hans-Georg Kräusslich, Roland Eils, Karl Rohr, Christoph Bräuchle, Barbara Müller, and Don C. Lamb. Dynamics of HIV-1 assembly and release. PLoS pathogens, 5:e1000652–e1000652, November 2009. 10.1371/journal.ppat.1000652.

18 Nolwenn Jouvenet, Sanford M. Simon, and Paul D. Bieniasz. Imaging the interaction of HIV-1 genomes and Gag during assembly of individual viral particles. Proceedings of the National Academy of Sciences of the United States of America, 106:19114–19119, November 2009. 10.1073/pnas.0907364106.

19 Jordan T. Becker and Nathan M. Sherer. Subcellular Localization of HIV-1 gag-pol mRNAs Regulates Sitesof VirionAssembly. Journalof virology, 91,March 2017. 10.1128/JVI.02315-16.

20 Luca Sardo, Steven C. Hatch, Jianbo Chen, Olga Nikolaitchik, Ryan C. Burdick, De Chen, Christopher J. Westlake, Stephen Lockett, Vinay K. Pathak, and Wei-Shau Hu. Dynamics of HIV-1 RNA Near the Plasma Membrane during Virus Assembly. Journal of Virology, 89 (21):10832–10840, November 2015. ISSN 1098-5514. 10.1128/JVI.01146-15.

21 Kari A. Dilley, Olga A. Nikolaitchik, Andrea Galli, Ryan C. Burdick, Louis Levine, Kelvin Li, Alan Rein, Vinay K. Pathak, and Wei-Shau Hu. Interactions Between HIV-1 Gag and Viral RNA Genome Enhance Virion Assembly. Journal of virology, May 2017. 10.1128/JVI.02319-16.

22 Jean-Baptiste Sibarita. High-density single-particle tracking: quantifying molecule organization and dynamics at the nanoscale. Histochemistry and Cell Biology, 141(6):587–595, June 2014. ISSN 1432-119X. 10.1007/s00418-014-1214-1.

23 Mohamed El Beheiry, Maxime Dahan, and Jean-Baptiste Masson. InferenceMAP: mapping of single-molecule dynamics with Bayesian inference. Nature methods, 12:594–595, July 2015. 10.1038/nmeth.3441.

24 Mohamed El Beheiry, Silvan Türkcan, Maximilian U. Richly, Antoine Triller, Antigone Alexandrou, Maxime Dahan, and Jean-Baptiste Masson. A Primer on the Bayesian Approach to High-Density Single-Molecule Trajectories Analysis. Biophysical journal, 110:1209–1215, March 2016. 10.1016/j.bpj.2016.01.018.

25 Jakub Chojnacki and Barbara Müller. Investigation of HIV-1 assembly and release using modern fluorescence imaging techniques. Traffic (Copenhagen, Denmark), 14(1):15–24, January 2013. ISSN 1600-0854. 10.1111/tra.12006.

26 Antony K. Chen, Prabuddha Sengupta, Kayoko Waki, Schuyler B. Van Engelenburg, Takahiro Ochiya, Sherimay D. Ablan, Eric O. Freed, and Jennifer Lippincott-Schwartz. MicroRNA binding to the HIV-1 Gag protein inhibits Gag assembly and virus production. Proceedings of the National Academy of Sciences of the United States of America, 111(26): E2676–2683, July 2014. ISSN 1091-6490. 10.1073/pnas.1408037111.

27 Audrey Thomas, Charlotte Mariani-Floderer, Maria Rosa López-Huertas, Nathalie Gros, Elise Hamard-Péron, Cyril Favard, Theophile Ohlmann, José Alcamí, and Delphine Muriaux. Involvement of the Rac1-IRSp53-Wave2-Arp2/3 Signaling Pathway in HIV-1 Gag Particle Release in CD4 T Cells. Journal of virology, 89:8162–8181, August 2015. 10.1128/JVI.00469-15.

28 A. Ono, D. Demirov, and E. O. Freed. Relationship between human immunodeficiency virus type 1 Gag multimerization and membrane binding. Journal of Virology, 74(11):5142–5150, June 2000. ISSN 0022-538X.

29 Jelle Hendrix, Viola Baumgärtel, Waldemar Schrimpf, Sergey Ivanchenko, Michelle A. Digman, Enrico Gratton, Hans-Georg Kräusslich, Barbara Müller, and Don C. Lamb. Live-cell observation of cytosolic HIV-1 assembly onset reveals RNA-interacting Gag oligomers. The Journal of cell biology, 210:629–646, August 2015. 10.1083/jcb.201504006.

30 Julia Gunzenhäuser, Nicolas Olivier, Thomas Pengo, and Suliana Manley. Quantitative super-resolution imaging reveals protein stoichiometry and nanoscale morphology of assembling HIV-Gag virions. Nano Letters, 12(9):4705–4710, September 2012. ISSN 1530-6992. 10.1021/nl3021076.

31 Maria R. D’Orsogna, Bingyu Zhao, Bijan Berenji, and Tom Chou. Combinatoric analysis of heterogeneous stochastic self-assembly. The Journal of Chemical Physics, 139(12): 121918, September 2013. ISSN 0021-9606, 1089-7690. 10.1063/1.4817202.

32 Rundong Hu, Baiping Ren, Mingzhen Zhang, Hong Chen, Yonglan Liu, Lingyun Liu, Xiong Gong, Binbo Jiang, Jie Ma, and Jie Zheng. Seed-Induced Heterogeneous Cross-Seeding Self-Assembly of Human and Rat Islet Polypeptides. ACS Omega, 2(3):784–792, March 2017. ISSN 2470-1343, 2470-1343. 10.1021/acsomega.6b00559.

33 Brian R. Novak, Edward J. Maginn, and Mark J. McCready. Comparison of heterogeneous and homogeneous bubble nucleation using molecular simulations. Physical Review B, 75 (8), February 2007. ISSN 1098-0121,1550-235X. 10.1103/PhysRevB.75.085413.

34 Pei-I. Ku, Anna K. Miller, Jeff Ballew, Virginie Sandrin, Frederick R. Adler, and Saveez Saffarian. Identification of pauses during formation of HIV-1 virus like particles. Biophysical Journal, 105(10):2262–2272, November 2013. ISSN 1542-0086. 10.1016/j.bpj.2013.09.047.

35 Bridget A Robinson, Jonathan C Reed, Clair D Geary, J Victor Swain, and Jaisri R Lingappa. A temporospatial map that defines specific steps at which critical surfaces inthe gag ma and ca domains act during immature hiv-1 capsid assembly in cells. Journal of virology, 88(10): 5718–5741,2014.

36 Nolwenn Jouvenet, Sanford M. Simon, and Paul D. Bieniasz. Visualizing HIV-1 assembly. Journal of molecular biology, 410:501–511, July 2011. 10.1016/j.jmb.2011.04.062.

37 Nicolas Chenouard, Ihor Smal, Fabrice de Chaumont, Martin Maška, Ivo F. Sbalzarini, Yuan-hao Gong, Janick Cardinale, Craig Carthel, Stefano Coraluppi, Mark Winter, Andrew R. Cohen, William J. Godinez, Karl Rohr, Yannis Kalaidzidis, Liang Liang, James Duncan, Hongying Shen, Yingke Xu, Klas E. G. Magnusson, Joakim Jaldén, Helen M. Blau, Perrine Paul-Gilloteaux, Philippe Roudot, Charles Kervrann, Frangois Waharte, Jean-Yves Tinevez, Spencer L. Shorte, Joost Willemse, Katherine Celler, Gilles P. van Wezel, Han-Wei Dan, Yuh-Show Tsai, Carlos Ortiz de Solórzano, Jean-Christophe Olivo-Marin, and Erik Meijer-ing. Objective comparison of particle tracking methods. Nature Methods, 11(3):281–289, March 2014. ISSN 1548-7105. 10.1038/nmeth.2808.

38 M. Chertkov, L. Kroc, F. Krzakala, M. Vergassola, and L. Zdeborová. Inference in particle tracking experiments by passing messages between images. Proceedings of the National Academy of Sciences of the United States of America, 107(17):7663–7668, April 2010. ISSN 1091-6490. 10.1073/pnas.0910994107.

39 Jean-Baptiste Masson, Patrice Dionne, Charlotte Salvatico, Marianne Renner, Christian G. Specht, Antoine Triller, and Maxime Dahan. Mapping the energy and diffusion landscapes of membrane proteins at the cell surface using high-density single-molecule imaging and Bayesian inference: application to the multiscale dynamics of glycine receptors in the neuronal membrane. Biophysical journal, 106:74–83, January 2014. 10.1016/j.bpj.2013.10.027.

40 Alexander J. Pak, John M. A. Grime, Prabuddha Sengupta, Antony K. Chen, Aleksander E. P. Durumeric, Anand Srivastava, Mark Yeager, John A. G. Briggs, Jennifer Lippincott-Schwartz, and Gregory A. Voth. Immature HIV-1 lattice assembly dynamics are regulated by scaffolding from nucleic acid and the plasma membrane. Proceedings of the National Academy of Sciences of the United States of America, 114(47):E10056–E10065, November 2017. ISSN 1091-6490. 10.1073/pnas.1706600114.

41 A. R. Evans, M. S. Turner, and P. Sens. Interactions between proteins bound to biomembranes. Physical Review. E, Statistical, Nonlinear, and Soft Matter Physics, 67(4 Pt 1): 041907, April 2003. ISSN 1539-3755. 10.1103/PhysRevE.67.041907.

42 Pierre Sens and Matthew S. Turner. Theoretical model for the formation of caveolae and similar membrane invaginations. Biophysical Journal, 86(4):2049–2057, April 2004. ISSN 0006-3495. 10.1016/S0006-3495(04)74266-6.

43 D. Muriaux, J. Mirro, D. Harvin, and A. Rein. RNA is a structural element in retrovirus particles. Proceedings of the National Academy of Sciences of the United States of America, 98(9):5246–5251, April 2001. ISSN 0027-8424. 10.1073/pnas.091000398.

44 Cendrine Faivre-Moskalenko, Julien Bernaud, Audrey Thomas, Kevin Tartour, Yvonne Beck, Maksym Iazykov, John Danial, Morgane Lourdin, Delphine Muriaux, and Martin Castelnovo. RNA Control of HIV-1 Particle Size Polydispersity. PLoS ONE, 9(1):e83874, January 2014. ISSN 1932-6203. 10.1371/joumal.pone.0083874.

45 Sebla B. Kutluay and Paul D. Bieniasz. Analysis of the initiating events in HIV-1 particle assembly and genome packaging. PLoS pathogens, 6(11):e1001200, November 2010. ISSN 1553-7374. 10.1371/joumal.ppat.1001200

46 Boyan Grigorov, Fabienne Arcanger, Philippe Roingeard, Jean-Luc Darlix, and Delphine Muriaux. Assembly of infectious HIV-1 in human epithelial and T-lymphoblastic cell lines. Journal of Molecular Biology, 359(4):848–862, June 2006. ISSN 0022-2836. 10.1016/j.jmb.2006.04.017.

47 Ignacio Izeddin, Christian G. Specht, Mickaël Lelek, Xavier Darzacq, Antoine Triller, Christophe Zimmer, and Maxime Dahan. Super-resolution dynamic imaging of dendritic spines using a low-affinity photoconvertible actin probe. PloS One, 6(1):e15611, January 2011. ISSN 1932-6203. 10.1371/joumal.pone.0015611.

48 Adel Kechkar, Deepak Nair, Mike Heilemann, Daniel Choquet, and Jean-Baptiste Sibarita. Real-time analysis and visualization for single-molecule based super-resolution microscopy. PloS One, 8(4):e62918, 2013. ISSN 1932-6203. 10.1371/joumal.pone.0062918

49 Arnauld Sergé, Nicolas Bertaux, Hervé Rigneault, and Didier Marguet. Dynamic multiple-target tracing to probe spatiotemporal cartography of cell membranes. Nature Methods, 5(8):687–694, August 2008. ISSN 1548-7105. 10.1038/nmeth.1233.

50 Hanns L. Harney. Form Invariance II: Natural x. In Bayesian Inference, pages 95–108. Springer Berlin Heidelberg, Berlin, Heidelberg, 2003. ISBN 978-3-642-05577-5 978-3-66206006-3. 10.1007/978-3-662-06006-3_11.

51 Fredrik Persson, Martin Lindén, Cecilia Unoson, and Johan Elf. Extracting intracellular diffusive states and transition rates from single-molecule tracking data. Nature Methods, 10(3):265–269, March 2013. ISSN 1548-7105. 10.1038/nmeth.2367.

52 Jan-Willem van de Meent, Jonathan E. Bronson, Chris H. Wiggins, and Ruben L. Gonzalez. Empirical Bayes methods enable advanced population-level analyses of single-molecule FRET experiments. Biophysical Journal, 106(6):1327–1337, March 2014. ISSN 1542-0086. 10.1016/j.bpj.2013.12.055.

53 Joshua C. Chang, Pak-Wing Fok, and Tom Chou. Bayesian Uncertainty Quantification for Bond Energies and Mobilities Using Path Integral Analysis. Biophysical Journal, 109(5): 966–974, September 2015. ISSN 1542-0086. 10.1016/j.bpj.2015.07.028.

